# Biophysical metabolic modeling of complex bacterial colony morphology

**DOI:** 10.1101/2024.03.13.584915

**Authors:** Ilija Dukovski, Lauren Golden, Jing Zhang, Melisa Osborne, Daniel Segrè, Kirill S. Korolev

## Abstract

Microbial colony growth is shaped by the physics of biomass propagation and nutrient diffusion, and by the metabolic reactions that organisms activate as a function of the surrounding environment. While microbial colonies have been explored using minimal models of growth and motility, full integration of biomass propagation and metabolism is still lacking. Here, building upon our framework for Computation of Microbial Ecosystems in Time and Space (COMETS), we combine dynamic flux balance modeling of metabolism with collective biomass propagation and demographic fluctuations to provide nuanced simulations of *E. coli* colonies. Simulations produced realistic colony morphology, consistent with our experiments. They characterize the transition between smooth and furcated colonies and the decay of genetic diversity. Furthermore, we demonstrate that under certain conditions, biomass can accumulate along “metabolic rings” that are reminiscent of coffee-stain rings, but have a completely different origin. Our approach is a key step towards predictive microbial ecosystems modeling.

## Introduction

Understanding the interplay of biological and physical processes in microbial growth is a long-standing endeavor of considerable relevance to multiple research areas.^1–7^ Microbial colonies on agar surfaces serve as a valuable model system to study ecological, evolutionary, and even developmental processes.^5,7–10^ Spatial and metabolic heterogeneity is ubiquitous across ecosystems,^11–13^ from complex communities^14,15^ of oceanic,^16–18^ or soil^19^ microbes to microbiota associated with plant roots,^20–22^ or animal digestive tracts^23^ to pathogenic communities growing on catheters^24^ and other medical implants. The familiar and apparently simple experimental system - *Escherichia coli* growing on an agar-filled Petri dish, is no exception. The growth of a microbial colony is a highly complex spatio-temporal phenomenon, involving the microenvironment-dependent metabolic activity and growth properties of the cells themselves,^8,25–27^ the diffusion of molecules and the propagation of cells on the agar,^28^ and the randomness induced by fluctuations in the birth-death process.^13,29,30^

Abundant work has been carried out to characterize experimentally biofilms on surfaces,^31,32^ showcasing a plethora of colony morphologies for different species and environmental conditions.^31,33–35^ Similarly, from a computational and theoretical perspective, there is a long tradition of developing reaction-diffusion^36–38,39,40^ and individual-based models^13,41^ to understand the origin of these spatial patterns. Such simple, phenomenological models have rationalized many empirical observations, but, given their simplicity, it is likely that some phenomena cannot be easily captured without a more mechanistic description. Moreover, phenomenological models do not explicitly cell physiology or growth conditions, and thus cannot be used to understand species differences or environmental drivers.

Despite their plain looks, microbial colonies are exquisitely complex and display distinct metabolic regimes in different microenvironments.^42^ Colonies grown on different media, and involving different organisms, could lead to drastically different behavior. This is due to the high dimensionality of parameters space involved: a typical bacterium harbors on the order of 10^3^ metabolites and reactions^43^; a typical agar plate involves dozens of nutrients; ^44^ and even the physical parameters of a Petri dish can play a key role in determining microbial physiology^1,3,36,45–48^. If one could simultaneously model all of these complexity, i.e. the metabolic activity of cells as a function of surrounding environment and the biophysical processes responsible for colony growth, it would be possible to achieve a predictive quantitative understanding of colony formation under varying conditions, with applications to biomedical research, environmental microbial ecology, and biotechnology.

Here, we merge the physical sophistication of the phenomenological models described above, with the biological accuracy and completeness of genome scale models, to implement a new biophysical model of microbial growth. Genome-scale models of metabolism, also known as stoichiometric constraint-based models, use information about the set of all known reactions in a cell and their stoichiometry to infer putative steady states that can maintain efficient growth given a set of environmental nutrients^43,49,50^. Previous efforts have taken important steps towards the development of integrated approaches in bacterial cellular^51,52^ and colony modeling,^53,54^ that combine the biophysical models of biomass propagation with the systems biology approach of genome-scale stoichiometric modeling^49^ of metabolism. However, to our knowledge, no single simulation framework has so far integrated genome scale modeling with the multiple biophysical processes known to potentially affect microbial growth in structured environments. We hypothesize that, in addition to providing a more comprehensive and flexible platform for biological systems simulation and quantitative predictive power, such an integrated framework would have the chance of uncovering new phenomena that depend on the concurrent effects of these multiple processes.

Our approach is to integrate the time-tested models of cellular biomass propagation,^55^ demographic fluctuations,^56,30^ and a model of core cellular metabolism,^50^ to produce remarkably realistic colony morphologies, and uncover novel colony features that depend on the interplay of physical and biological processes (Fig. 1). At the core of our method is the genome scale modeling of metabolism, as implemented in our software platform COMETS (**C**omputation of **M**icrobial **E**cosystems in **T**ime and **S**pace).^54^ This mechanistic description of cellular metabolism makes it possible to obtain growth rates and nutrient fluxes in an environment-dependent way, alleviating the need to restructure the model upon changing conditions and creating a direct correspondence between the time scales in the experiments and simulations. Metabolic fluxes are determined dynamically across space via dynamic Flux Balance Analysis (dFBA)^27,57^ carried out independently for each point of a spatial grid.

**Figure 1.**
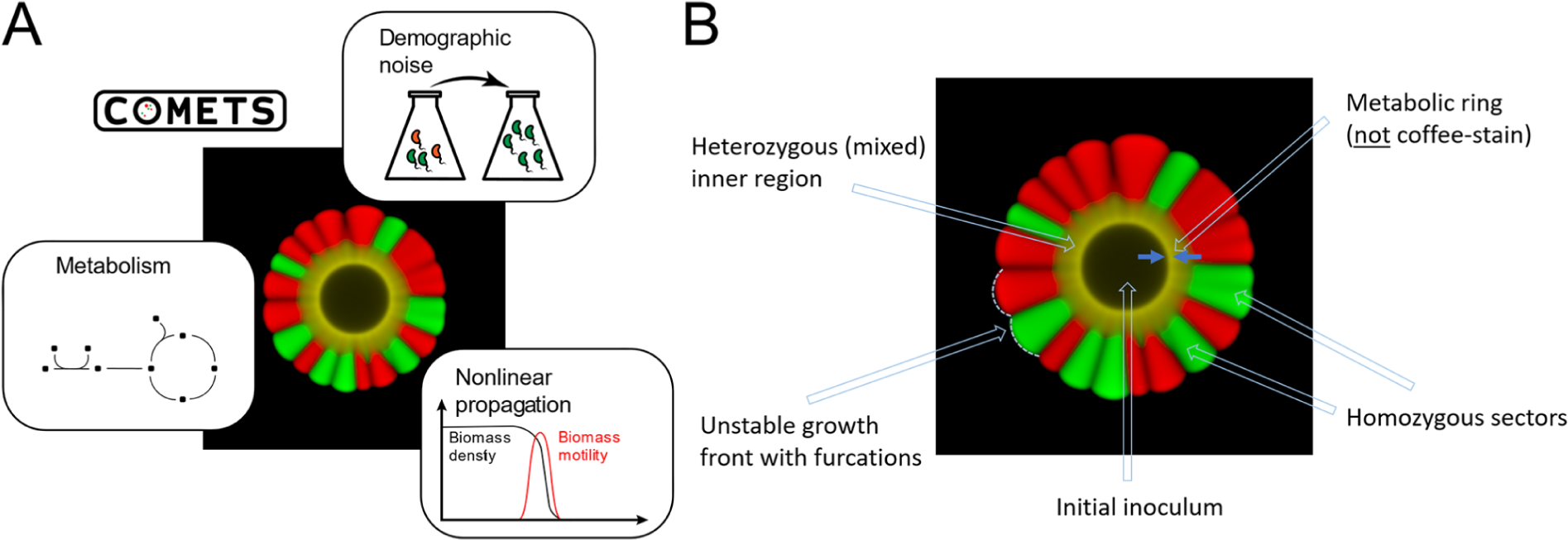
The integrated simulation methodology of COMETS realistically reproduces several colony features. (A) Schematic overview of the three methods implemented in COMETS: (i) the genome-scale stoichiometric modeling of the cellular metabolism, (ii) the nonlinear biomass propagation model, and (iii) the demographic noise. (B) Simulations capture some salient features of a bacterial colony: (i) initial inoculum surrounded by a metabolic ring (not related to the coffee-stain effect), (ii) heterozygous region (yellow) during early stages of colony growth, (iii) single strain homozygous regions (red and green), and (iv) unstable branching (furcating) growth front.

Previous versions of COMETS were handling biomass propagation through standard linear diffusion equations.^27^ This approach has been implemented also in other frameworks for modeling microbial growth, leading to valuable insight.^53,58–60^ However it is known that the propagation of cells through a biofilm is strongly affected by additional processes, which we added in our newly released version of COMETS. A first aspect we now explicitly take into account is the presence of cooperative and nonlinear processes, which can greatly affect colony morphology. In particular, it is known that microbial biomass can behave in ways that are drastically different from inanimate matter, most notably through biophysical processes that cause cells to diffuse more efficiently when surrounded by many other cells.^61–63^ This cooperative propagation of cellular biomass can be caused by a number of distinct phenomena, including mechanical interactions (cell-cell pushing), production of a polymer matrix, or secretion of a carrier fluid that enables swimming, *etc*. In all such situations, the higher the biomass density, the greater the cumulative effect of their collective propagation, enabling more efficient motion of individual cells.

A second novel aspect of our model is the implementation of demographic fluctuations^56,30^ due to the stochastic nature of the cellular birth and death processes. The fluctuations create population irregularities which could be damped or amplified by the growth dynamics. Perhaps, more importantly, demographics underlie all evolutionary modeling and can even cause local extinctions and fixations of a specific genotype.^64^ In the context of colony growth, these fixation events manifest in the formation of monoclonal sectors,^12,29,62,65^ which are typically visualized with fluorescent markers (Fig. 1B).

As described below, our new COMETS enables the simulation of many emergent properties of biomass propagation on surfaces. We first recapitulate previous findings in minimal models, but then extend them to complex metabolic models and new environmental conditions. We also report fundamentally new phenomena such as the emergence of a “metabolic ring” that is visually similar, but conceptually very different from the classical coffee stain ring caused by evaporation-induced fluid flows.^66–69^ The new version of COMETS paves the way for better integration of biological and physical processes, with important consequences for predictive modeling of biological systems in structured environments.

## Results

### Detailed biophysical models predict colony size and expansion rate across environments

We created a new version of COMETS that combines the metabolic dFBA components described before^54^ with a new more realistic description of the biophysical processes responsible for biomass propagation. The implementation of dFBA, unchanged relative to prior versions, solves FBA in iterative discrete time steps, by translating extracellular metabolic concentrations into uptake flux bounds, and effectively assuming that intracellular metabolism reaches steady state faster than the rate at which external resources are accumulated or depleted. While the mathematical details of the approach and a description of the computational implementation are described thoroughly in the Methods section, we highlight here briefly the major achievements of the newly designed COMETS algorithm.

The points in the COMETS spatial grid are assumed to exchange with each other both metabolites via regular diffusion and biomass via a propagation process. The latter is described by a biophysical model of cooperative biomass propagation.^13,55^ The cooperative mode of biomass propagation emerges in growth conditions where cells cannot move independently, e.g. by swimming and instead rely on other mechanisms such as pushing onto each other as they grow. The spreading of the biomass in such crowded conditions is often cooperative since the dense the packing the more effective the pushing. Hence, in our model, the effective diffusivity of the biomass (*D_b_*) increases with the biomass density (b) as: *D_b_* = *D*_0_*b^k^*, where *D*_0_ and *k* are parameters of the model. Note that here we purposefully neglect the presence of additional diffusion terms associated with active motility (see also Discussion). Based on prior evidence, ^13,55^ we assume that *k* = 1 throughout all the simulations presented here. It is also important to note that the capacity of cells to propagate on the surface (*D*_0_) is growth-dependent (see Methods). This assumption is based on experimental observations of reduced cell-motility in the colony center due to nutrient limitation, extracellular matrix or mechanical jamming.^70^

The new version of COMETS also includes an implementation of demographic fluctuations, which affect both the overall shape of the colony and the spatial distribution of genetic variants (sectors).^12,13,29^ The variance of demographic fluctuations is proportional to the local biomass concentration, and their probability distribution is approximately Gaussian except at very low biomass densities.^29^ We relied on this Gaussian approximation, but ensured that the biomass remains non-negative. This was achieved by deploying the split-step method which separates the deterministic from the stochastic simulation step^56^ (see Methods). In some simulations, we also included non-demographic, e.g. environmental, fluctuations by adding a normally distributed random variable to the growth rate predicted by dFBA (see Methods).

To illustrate the value of a model with a detailed metabolic network, we compared it to a minimal phenomenological model while keeping everything else equal. The stoichiometry-based model, different sources of carbon, nitrogen and other nutrients can be absorbed, metabolized, interconverted and ultimately used to synthesize biomass with specific stoichiometric ratios as described in the standard model of core *E. coli* metabolism.^50^ The minimal model (based on prior work by Müller and Van Saarloos (2002)) can also be implemented in COMETS by defining a single reaction that directly converts a growth-limiting nutrient (in this case, glucose) into biomass (see Methods). To make a fair comparison, all non-metabolic aspects such as biomass propagation and demographic noise were identical for the two models. The only difference between the simulations was in the cellular metabolic networks. The single-nutrient model contained a single reaction that converted glucose into biomass. We tested the single-nutrient model at two levels of biomass to glucose yield ratios. First we simulated a yield ratio of one, as implemented in prior theoretical work.^55^ Next we adjusted the glucose to biomass ratio to the mean batch culture yield of the *E. coli* core model (see Methods). For the metabolically realistic COMETS, we used a model of the core metabolism of *E. coli*^50^ (see Methods). This model has been extensively studied, and the uptake parameters reproduce growth rates of *E. coli* in experiments.^57^

We started with a growth medium that contained glucose as the only growth-limiting nutrient. The minimal model (with yield=1) and the COMETS *E. coli* models produced very different results (Fig. 2A and 2B) both in terms of time-scale, but also in terms of morphology features. While the COMETS model tracked realistically the typical growth curve of *E. coli*,^26,71^ the minimal model predicted a much higher growth rate presumably due to the unrealistic yield, and to the fact that it did not account for the various “overhead” reactions necessary to build a cell. In part because of this growth rate difference, the morphologies in the two simulations were also distinct. The minimal model predicted a nearly smooth while the colony had strong front furcations in the COMETS simulation.

**Figure 2.**
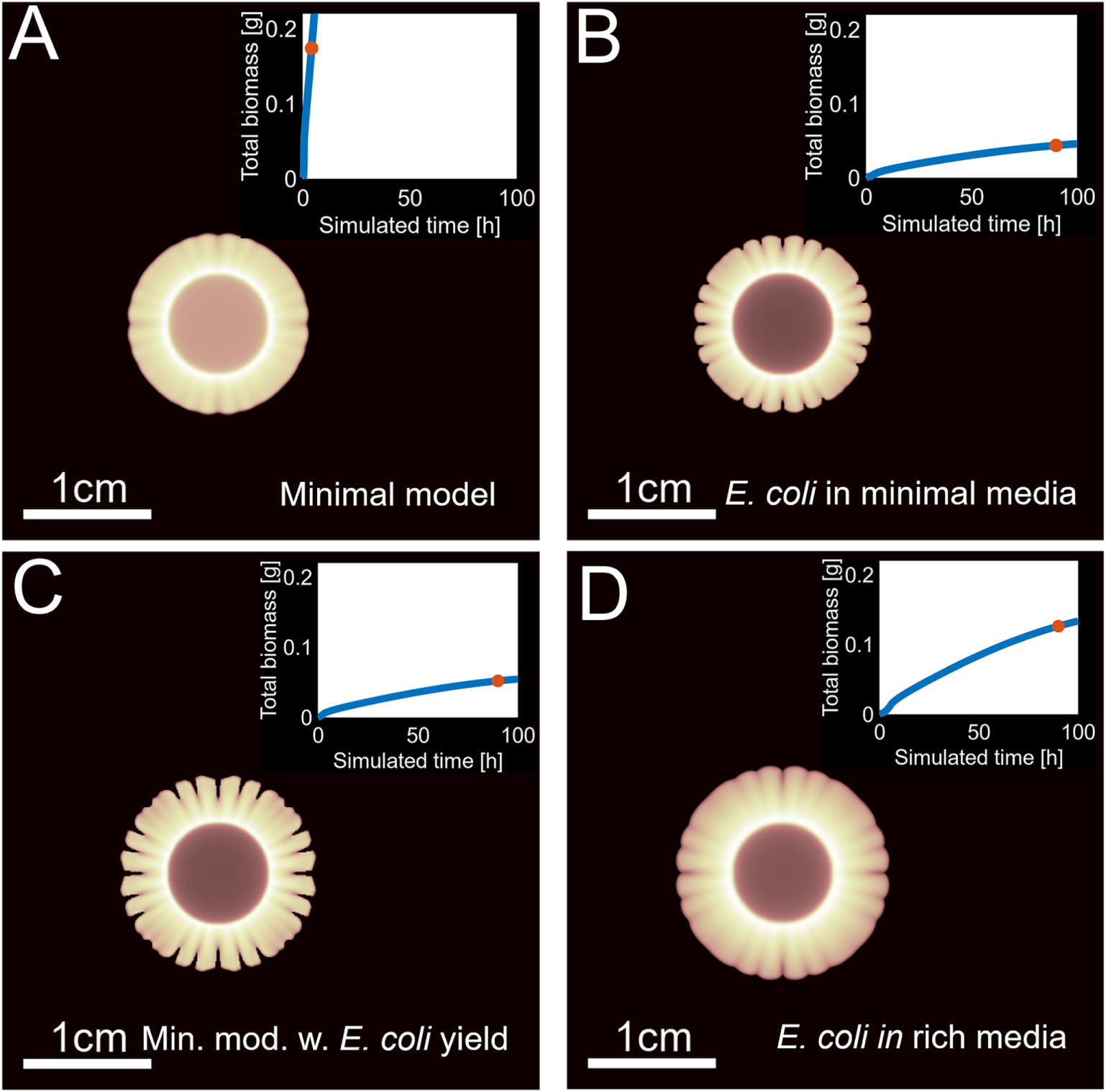
Genome-scale metabolic modeling is more realistic than the single-metabolite minimal model. Examples of simulated bacterial colonies modeled with: (A) Minimal model in presence of a single nutrient. The entire metabolic network consists of a single reaction that converts the nutrient into biomass. (B) *E. coli* core model in a complex nutrient substrate with glucose as a limiting carbon source; similar to a real bacterium, the model cannot grow on just glucose and needs minerals, cofactors, and a source of nitrogen. (C) the minimal model as in (A), with the biomass yield set to produce the same amount of biomass from a single nutrient as the amount produced by the *E. coli* stoichiometric model fed an equivalent amount of glucose. (D) *E. coli* stoichiometric model in complex nutrient substrate similar to the substrate in (B), but with ammonia replaced with glutamate as a source of nitrogen. The biophysical model for spatial propagation of the biomass is exactly the same in all three cases. The insets show the growth curves and the red dots refer to the time points for which the colony images are shown. Note that COMETS predicts colonies comparable in size with those observed experimentally, while the minimal model in (A) predicts a much faster growth, unless its yield is adjusted to match the *E. coli* model in (C).

When we adjusted the biomass to glucose yield ratio in the minimal model to the one that matches the yield of the *E. coli* core model batch culture (Fig. 2C), the growth curve and the colony morphology came much closer (although not exactly the same) to the one produced by the *E. coli* core model (Fig. 2B). Thus the minimal model, with appropriate manual tuning of the biomass yield or the nutrient concentration, seems to approximate reasonably well the much more complex COMETS model. The limited utility of such a fitting procedure, however, becomes apparent when environmental conditions are changed. As an example of this, we repeated the simulations described above with the same glucose concentration, but with ammonia replaced by glutamate as a source of nitrogen (see Fig. 2D). This substitution effectively replaces the minimal medium in Fig. 2B by a richer medium in Fig. 2D. Consistent with experiments, the simulations predicted much faster growth in the rich media. It would be impossible to make such a prediction using the minimal model. One would have to artificially adjust either the concentration of the carbon source or the biomass/carbon source yield to artificially match the observations.

Not only the minimal model requires a different fit for any change in composition of the growth media, but also it is completely incapable of describing diauxic shifts^72,49,57^ or other changes in nutrient consumption during colony growth. Similarly, the minimal model would not be able to take into account metabolic regime changes that may be induced by local depletion of metabolites or accumulation of byproducts, let alone cross-feeding in a multi-species consortium. Thus, a realistic description of cellular metabolism is essential if one hopes to capture the whole spectrum of growth patterns that can occur in any meaningful experiment.

### COMETS simulations for E. coli match experimentally-observed growth curves and colony morphologies

To assess the accuracy of COMETS predictions, we compared the growth of a simulated *E. coli* colony to experiments performed under the same conditions (Fig. 3). The main plot in Fig. 3 displays the simulated colony radius, compared with corresponding experimental measurements (see also growth curve of the total (integrated) colony biomass in the inset). The growth is initially close to linear, consistent with prior theoretical and experimental studies.^73^ As nutrients become scarce, growth slows down (see Fig. 3, inset) and eventually stops around day six. The experiments were carried out in a six-well plate, on relatively hard agar with R2A media supplemented with glucose (see Methods). The simulations used exactly the same spatial dimensions. In simulations, the nutrient concentrations were adjusted to account for the differences between the composition of the R2A media and the metabolites included in the *E. coli* core model (see Methods). The three fitting parameters were the biomass diffusivity, the diffusivity cutoff (see Methods) and the strength of the demographic noise, of which only the biomass diffusivity plays a major role in this case.

**Figure 3.**
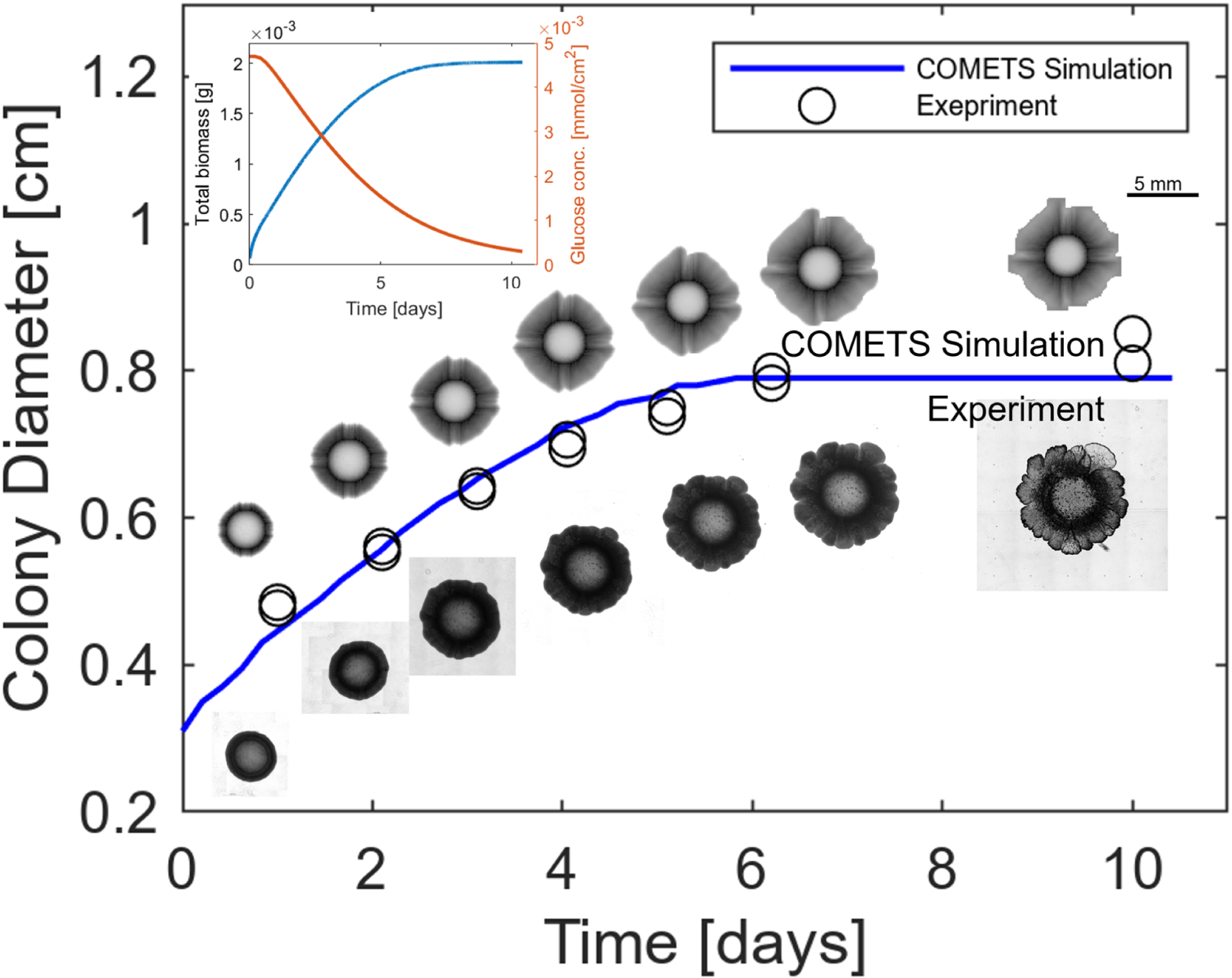
COMETS simulations of the growth of an *E. coli* colony are very close to the actual colony growth observed experimentally. The main curve shows the diameter of the simulated colony. Below the growth curve are 4x microscopy images (taken daily) of an *E. coli* colony grown on an agar plate in the lab. The images above the growth curve show the corresponding snapshots of the simulated *E. coli* model. The inset shows the total simulated biomass, and the simulated depletion of glucose at a single spatial point at the plate edge, 1.5 cm from the center.

In addition to the quantitative comparison of the colony radius, it is useful to perform a qualitative comparison of colony morphologies. The agreement between the experiments and simulations can be judged from the images of actual (microscopy, see Methods) and simulated (COMETS) colonies shown below and above the growth curve respectively. We see that both actual and simulated colonies begin to grow slower around day 6, have similar roughness of the colony edge, and exhibit a comparable accumulation of the biomass around the inoculation site (see the last section in Results).

### Demographic noise and biomass motility control branching and sectoring

The visual agreement between experiments and simulations encouraged us to ask whether we can successfully reproduce not only the temporal dynamics of the biomass growth, but also the spatial morphology along all of the stages of the colony development under different conditions. Specifically, a question that has attracted a lot of interest in the past and has been studied both experimentally^46,47^ and with minimal models^13,60^ is how colony shape is influenced by the agar hardness and the concentration of the nutrients in the substrate. Characteristic colony morphologies were found in different regions of the agar-nutrient “phase space” for *E. coli* ^47^ and other bacteria^45^ (see Fig. 5).

As a first step, we sought to recapitulate the most salient changes in colony morphology, i.e. the transition between smooth and rough (or furcated) fronts. In both experiments and minimal models, furcated fronts were observed under reduced cellular motility or nutrient concentration. The onset of front roughness is driven by the shifting balance between a stabilizing effect of biomass motility and a destabilizing effect of nutrient diffusion. Whenever a small protrusion appears, biomass diffusion tends to even it out. In contrast, nutrient diffusion leads to a greater nutrient flux towards the tip of the protrusion thereby increasing its growth rate and further destabilizing the interface.

We wanted to confirm whether a similar transition between smooth and furcated fronts can be observed in COMETS. To that end, we varied the biomass motility by adjusting *D*_0_ in the model, while keeping all other parameters including the nutrient diffusivity fixed. We indeed found a transition between smooth and rough fronts (see Fig. 4A). To quantify the transition we used the standard measure of interface roughness, defined as the root mean square deviation of the interface “height” (outward expansion distance, the y-coordinate) averaged over the length of the front (the x-coordinate). We found that the roughness stayed constant at high values of *D*_0_; in this regime front undulations reflect demographic fluctuations and are expected to fall into Kardar-Parisi-Zhang (KPZ) universality class.^74^ Below a certain threshold, however, roughness increased rapidly as *D*_0_ became smaller, indicating the onset of a growth instability.

**Figure 4.**
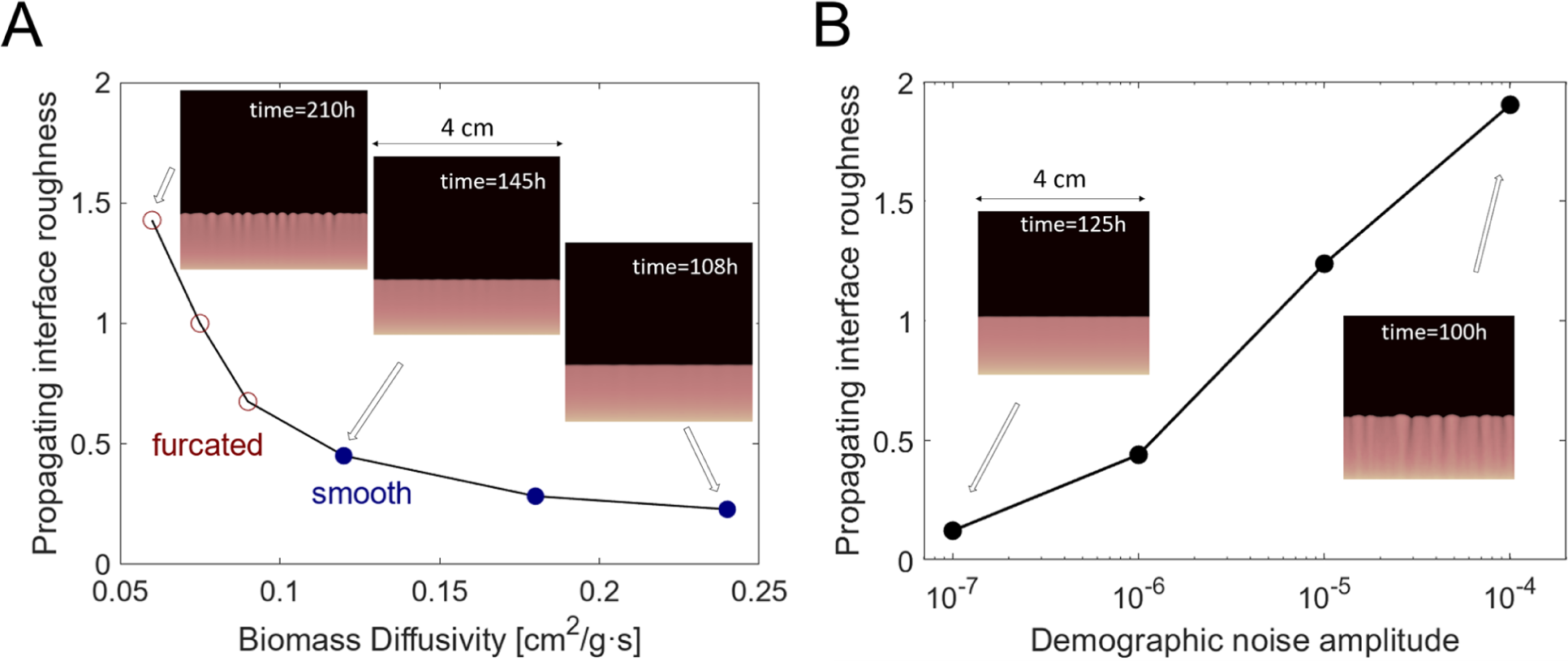
The transition from furcated to smooth growing front for the *E. coli* model. (A) The morphology of the growth front changes as the biomass diffusivity is increased. For low biomass diffusivity, the front is furcated (red empty points), but the front is smooth (solid blue points) at higher biomass diffusivities. The fronts are never perfectly flat because of the demographic fluctuations (*σ* = 10^−7^) and the additional noise in the growth rate (about 0.3%). To distinguish the regime of instability (red circles) from the regime of noise-induced roughening (blue dots) we examined the peak of the Fourier transform of the growth front position as shown in Fig. S1. The peak in the transform decays across the transition from furcated to smooth fronts, but then remains constant as the biomass diffusivity increases even further. (B) Higher amplitudes of the demographic noise lead to a more furcated growth front. All simulations started with a flat uniform biomass at the lower edge of the square layout. The diffusivity of the nutrients was the same 6·10^-6^ cm^2^/s in all simulations. The nutrients were initially supplied in the entire layout, and subsequently kept at that concentration at the top edge of the layout.

**Figure 5.**
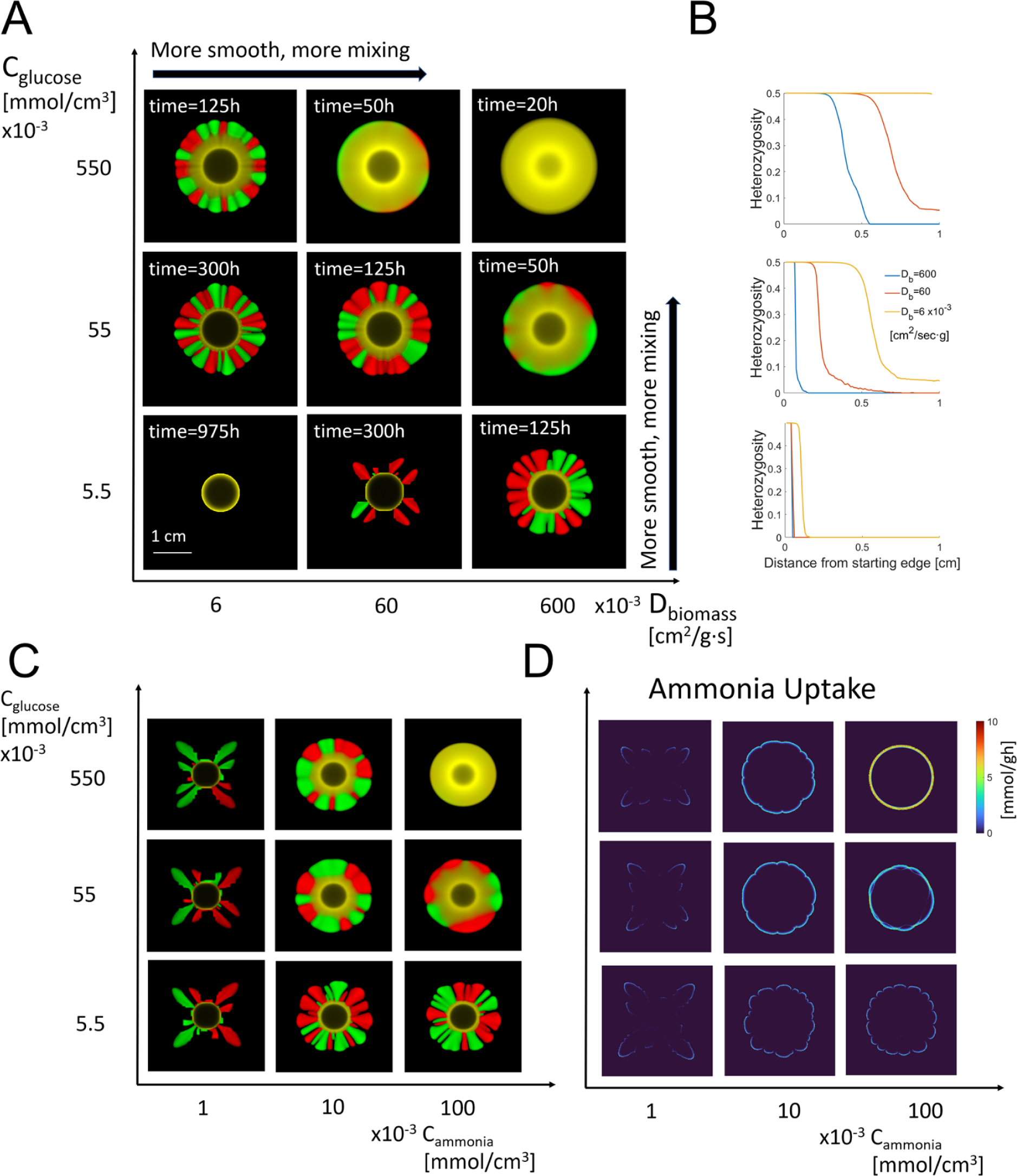
Genetic demixing and colony morphology show a similar pattern of variation across different growth conditions for the *E. coli* model. (A) Snapshots of fully-developed colonies for varying nutrient concentration and biomass motility. Note that both factors promote growth, reduce branching, and suppress genetic demixing. Different colors represent two metabolically-identical strains of *E. coli*. (B) Quantitative characterization of genetic demixing, in the direction of the colony growth. Here, heterozygosity is the probability to sample two different colors at the same spatial location. (C) The colony morphology is determined by the concentration of the limiting nutrient. Here we varied the concentration of glucose as a source of carbon, and ammonia as a source of nitrogen. For very low values of ammonia, the change of glucose concentration has no effect on the colony morphology. The growth of the colony is dictated by the uptake of ammonia, as illustrated in (D). These simulations were carried out in a square 4cm by 4cm spatial domain with a fixed nutrient concentration at the boundary. These boundary conditions allowed us to delay nutrient depletion and thus approximate growth in a larger Petri dish. Initially, the nutrient concentration was uniform and matched the boundary conditions. The diffusivity of the nutrients was 6·10^-6^ cm^2^/s across all conditions. We used minimal media with minimal salts and glucose as a source of carbon. The colonies were inoculated with a “flat” homeland drop at the center of the simulation domain (see Methods).

While, historically, the above transition was studied deterministically, we took advantage of our new platform to explore the effects of demographic noise on this transition (Fig. 4B). We simulated colony at a fixed value of the biomass diffusivity (*D*_0_ = 0.21*sm*^2^/*g* · *s*), a value moderately removed from the onset of the growth instability (see Fig. 4A), and increased the demographic noise amplitude over four orders of magnitude. The interface roughness increased with noise and eventually approached the values found for unstable growth at low *D*_0_. This increase however was very slow and without a sharp transition. Therefore, we conclude that demographic fluctuations facilitate the onset of branching by promoting the growth instability and should be included not only in simulations of colony growth but also in related models of tissue invasion by pathogens and even tumor growth.

Given that our simulations fully recapitulate the classic transition from smooth to furcated fronts, we can now explore new questions about the morphology and diversity of microbial colonies. To that end, we carried out simulations of initially circular colonies, which is the typical shape after inoculating with a drop of a liquid culture. We also included genetic diversity to capture the formation of sectors. This was accomplished by starting the simulations with a homogeneous mixture of two neutral genotypes, i.e. strains with identical metabolic networks and biomass diffusivity, but a distinguishing heritable marker such as a chromosomal insertion of a gene encoding a fluorescent protein (Fig. 5A).

Note that, since these neutral strains have identical growth rates, the colony morphology is not affected by the fact that we start with a mixture. Below, we first describe colony morphologies and then proceed to discuss sectoring.

Biomass diffusivity *D*_0_, as we saw in Fig. 4, is only one of many parameters that can drive the transition between smooth and furcated fronts. Historically, most of the attention has been paid to the nutrient availability and biomass diffusivity because these can be easily tuned in the experiments by varying glucose and agar concentrations respectively. The corresponding phase diagram for our *E. coli* model recapitulates previous theoretical and experimental findings and is shown in Fig. 5A. Little to no growth occurred at low nutrient concentration and biomass diffusivity (bottom left corner in Fig. 5A). Moreover, most of the growth occurred near the colony edge resulting in a ring-like appearance of the final colony (see the next section). A branched morphology was observed when either nutrient concentration or biomass diffusivity were increased. An increase in the biomass diffusivity produced more disjoint branches compared to an equal increase in the glucose concentration. The branches tended to thicken and merge as the growth conditions became more favorable. At the top right corner of the diagram, we found another morphology—a smooth disk. This is consistent with empirical observations of smooth round colonies found in soft agar with excess nutrients.^47^

Our simulations also provide evidence of the importance of metabolic details for an accurate description of colony growth. For example, we found that the increase in the glucose concentration had a stronger effect on the colony morphology than the same-fold increase in the biomass diffusivity. Previous work on minimal models suggested that the two parameters are interchangeable and, in fact, all aspects of colony morphology are controlled by a single dimensionless parameter proportional to *C_glucose_* · *D*_0_.^13^ While this prediction holds largely true in our simulations, there are some noticeable departures as well. These can be seen clearly along the main diagonal, which shows different branching patterns for three conditions with identical *C_glucose_* · *D*_0_. Our results thus show that some predictions of the minimal models rely on their simplifying assumptions about the functional form of the growth rate and do not hold in general.

To better understand the differences between the minimal and the *E. coli* models, we varied nutrient uptake parameters in our simulations. In the minimal model, the nutrient uptake is linear in both the nutrient concentration and the biomass density. A more general Michaelis-Menten type kinetics is used in COMETS (see Methods). We first examined how colony morphologies depend on the the maximum uptake parameter *V_max_* (Fig. S4). A greater value of the maximum uptake leads to a greater growth rate, which of course changes the relative time scales of growth, motility, and nutrient diffusion. As a result, the morphologies at different *V_max_* were quite different regardless of whether we compared colonies of the same total biomass (Fig. S4a) or the same radii (Fig. S4b).

We then examined the effects of varying the Michaelis constant *K_M_* (Fig. S5). The colony morphologies remained virtually unchanged when *K* was varied by two orders of magnitude around its value for *E. coli* growing on glucose. This lack of sensitivity can be easily explained by the fact that the value of the Michaelis constant is very low compared to the limiting nutrient concentrations below which the growth stops, so the growth is predominantly occurs at very high, close to maximum, uptake rates, governed by *V_max_* and not by *K_M_*. Thus, unlike in the minimal model, the nutrient uptake in COMETS was proportional to the biomass but not the nutrient concentration. Despite this major difference, the branching morphologies obtained in both types of simulations were remarkably similar, which implies that the functional form of nutrient uptake is not the major controlling factor of colony shapes. Note that these conclusions may not extend to a different organism growing in a different environment, which again highlights the utility and ease-of-use of simulations grounded in realistic biochemistry compared to the minimal model.

Colony morphology is only one manifestation of the spatio-temporal growth process in microbial population dynamics. To get a better picture of how environmental conditions affect population dynamics, we looked into the changes of genetic diversity during colony growth. Our simulations allow for inclusion of several types that have identical growth rates and can be viewed as neutral mutants or fluorescently labeled strains. Because the mutants are neutral, the colony growth and morphology is not affected by whether the simulations are started with a single strain or a mixture. The opposite, however, is not true: the growth dynamics of the colony and its morphology have a strong effect on genetic drift, i.e. on the spatial distribution and temporal changes of the abundances of the two strains.

To probe these evolutionary dynamics, we started simulations with a uniform distribution of the two strains within the inoculation site. Because the strains are labeled as green and red, the mixed region appears as yellow (see Fig. 5A). As the colony grew, it either stayed mixed, e.g. at high glucose concentration and high biomass motility, or the two strains segregated spatially forming red and green sectors. This genetic demixing reflects local extinction of one of the two strains due to demographic fluctuations. We emphasize that the strength of the demographic noise was the same across all the environments shown in Fig. 5A. Thus, the different rates of extinctions reflect the difference in the degree to which the growth dynamics of the colony amplify or suppress demographic fluctuations.

Similar to previous studies,^13^ we find that sector formation and branched morphologies go hand in hand. In all branched colonies, each of the branches is homozygous for a particular color. This fixation occurs in the early stages of colony growth at low nutrient concentrations. In contrast, a clear region of mixed growth (yellow ring) is present at higher nutrient concentration (and to a lesser extent at higher biomass motility). Under the most favorable growth conditions, genetic demixing is weak or completely absent.

The same conclusions can be drawn from Fig. 5B, which shows how the probability to sample two different colors at the same spatial location depends on the radial distance away from the inoculation site. While this metric, known as local heterozygosity, loses the information about the sector shapes, it removes any ambiguity in interpreting the color changes in Fig. 5a. In particular, the quantitative plots in Fig. 5B shows that the demixing transition is rather sharp, and there are significant regions of the colony which are either fully mixed or fully demixed.

One of the advantages of including the full stoichiometric metabolic network in our model of *E. coli* is that we can study the role of nutrients other than the carbon source on the morphology of a colony. If carbon is plentifully supplied, nitrogen or phosphorus supply may be what limits the growth of the biomass, and that limiting factor may affect the spatial spread of the colony, and consequently its morphology [ref 69]. To probe such dynamics, we examined the role of ammonia as a limiting source of nitrogen.

Figure 5C shows the effects of glucose and ammonia supply. Regardless of the glucose concentration, when ammonia concentration is low, colonies consist of a few spread-out branches fixed for one of the two genotypes. This is similar to the effect of glucose limitation in Fig. 4A, which suggests that colony dynamics, albeit not exactly the same, are similar when some of the required nutrients are scarce. When ammonia is abundant, the colony growth is controlled by the glucose concentration, and we recover the results reported if Fig. 4A. At intermediate concentrations of ammonia, both nutrients have comparable effect on colony growth broadly consistent with the idea that higher nutrient levels result in fewer branches and slower genetic demixing.

These conclusions are further supported by Fig. 5C, which shows ammonia uptake rates across nutrient concentrations. When one of the nutrients is scarce the uptake rate is low and limited to the tips of the growing branches. Note that abundance of the other nutrient cannot compensate, i.e. the uptake rates remain low despite either ammonia or glucose being in excess. Only when the concentration of both nutrients is increased do we observe a corresponding increase in the ammonia uptake rates and the concomitant change in the roughness of the colony edge.

### Initial colony growth results in a “metabolic ring” around the inoculation site

Most of the experimental and theoretical work presented here so far focused on the two-dimensional shapes of microbial colonies. The colony images in Fig. 3 however, point to a strong variation in the biomass density along the radial direction. Since cells are nearly incompressible, density variations likely reflect also the changes in colony thickness. Thus, our simulations can describe colony shape in all three spatial dimensions.

Perhaps the most striking three-dimensional feature in our simulations is a prominent circularly-symmetric peak of cell density near the inoculation site. At first sight, this ring of excess biomass resembles the coffee-stain or coffee ring effect, i.e. the pileup of cells along the boundary of the initial droplet placed on the Petri-dish. The coffee-ring effect has been documented in both living ^67^ and nonliving^66,69^ matter, and it occurs due to boundary pinning and outward fluid flow during evaporation.^66^ Evaporation typically lasts less than an hour, i.e. the coffee ring is formed before the colony starts to expand. Our simulations, however, do not account for these aspects of pre-growth fluid dynamics and are initialized with a flat, pancake-like mini colony. Thus, the biomass ring in our simulation is due to the growth dynamics itself, rather than to not the classical coffee-ring effect. To make this distinction clear, we will refer to the ring emerging in our simulations as a “metabolic ring”.

The emergence of the metabolic ring is shown in Figure 6A and Video 1. The simulations begin without a coffee-ring (pancake-like initial conditions), but the metabolic ring quickly appears and is clearly visible at 10h. The difference between the metabolic ring and the growth front is, however, not apparent at this time. The two regions, however, become distinct as the colony expands. In particular, the last time point (100h) shows that the colony consists of three regions: a sharp increase in biomass density at the edge, the more gradual increase in thickness from the edge towards the metabolic ring, and a low density region in the colony interior (FIg. 6A).

**Figure 6.**
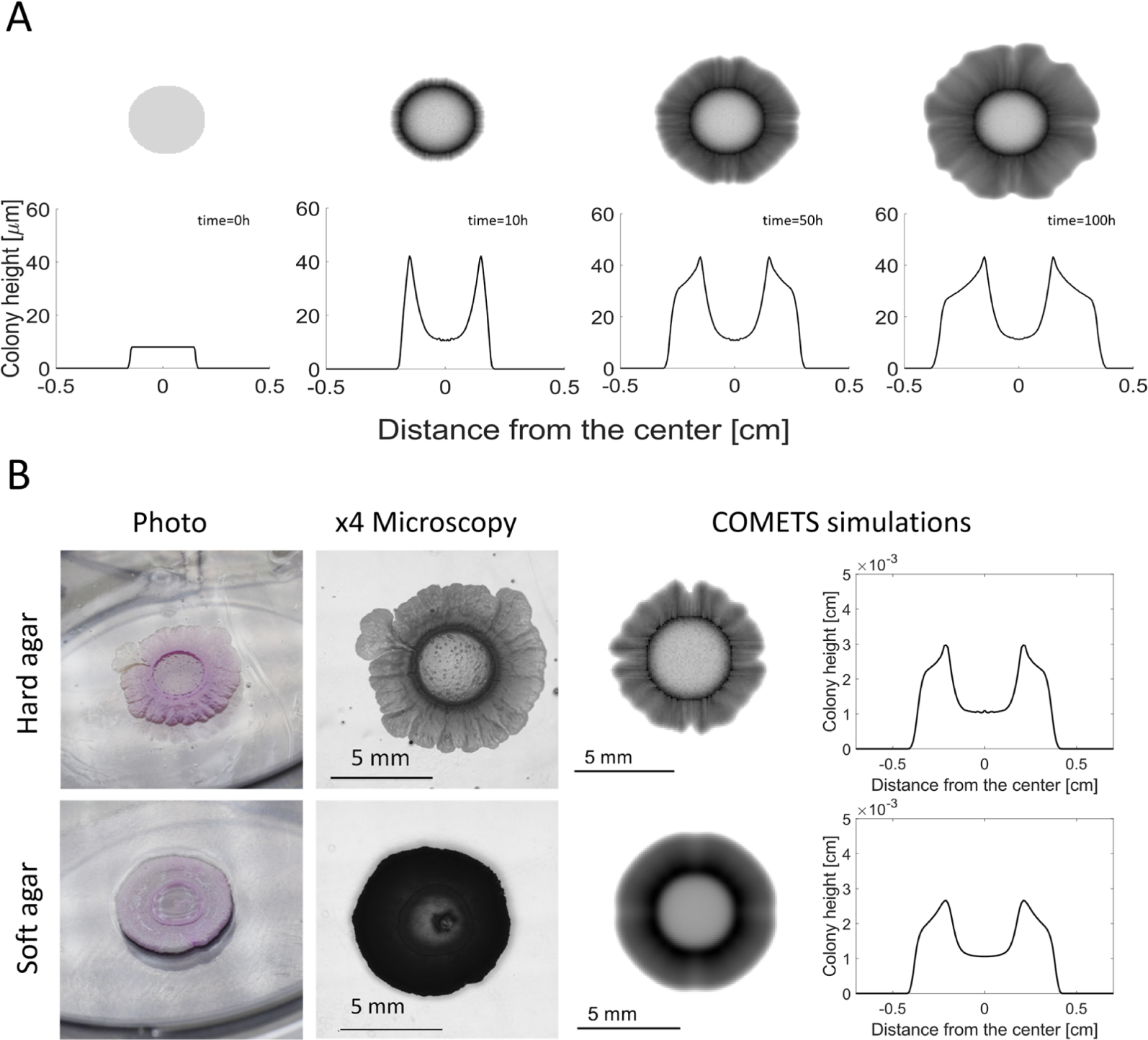
COMETS simulations predict a ridge of higher biomass growth around the initial colony homeland. The COMETS biophysical model does not include any capillary effects, which makes it possible to distinguish the metabolic ring from the similar capillary coffee-stain effect. (A) A growth sequence demonstrating the formation of the metabolic ring. The top images are simulated colonies viewed from above. The bottom images are estimated colony heights based on the biomass content in each pixel of the simulation grid, assuming that the biomass density is the same as that of water. Note that the ring forms very early, right before the colony starts to expand outward. See also Supplemental Video. (B) The metabolic ring is more prominent for colonies grown on hard agar (top row) than on soft agar (bottom row) in both experiments and simulations. The concentration of all nutrients is exactly the same and the only difference between the two growth conditions is the agar concentration in the experiments and biomass motility and demographic fluctuations in the simulations. The latter was adjusted to better match the roughness of the colony front, and this adjustment might reflect the real effect of harder agar on the strength of demographic fluctuations.

Our simulations predict that the biomass content of the metabolic ring (a proxy for colony thickness) varies with the environmental conditions. The ring is most prominent at low nutrient concentrations and low biomass diffusivity, but it is much weaker in more favorable growth conditions. In order to test this prediction, we grew *E. coli* colonies on substrates with several different agar concentrations (see Methods and Fig. S4). We highlight some of these data in Fig. 6B, which shows that the ring around the inoculation site is more prominent on hard agar in both simulations and experiments.

This agreement suggests that the capillary coffee-ring effect cannot be the sole explanation for the ring-like accumulation of the biomass near the inoculation site. Indeed, there is no a priori reason to expect the strength of the capillary coffee-ring effect to depend on the agar concentrations.^75^ Consistent with this reasoning, the early stages of colony growth show a clear coffee-ring effect for all agar concentrations (Fig. S4), but this ring continues to grow only in the colonies on hard agar as expected from our simulations.

A direct way to confirm the primary role of the growth dynamics in the formation of the biomass ring would be to eliminate the capillary coffee-ring in the experiments. While we believe that this is possible in principle, it is likely to be a difficult experiment.^69,76^ Instead, we used initial conditions in our simulations to mimic the presence or absence of the coffee ring and thus untangle their contribution to the final colony shape.

Specifically, we used three types of initial conditions: (i) a spherical drop with the maximal density in the center and zero density at the edge, (ii) a pancake with a uniform density within the inoculation disk, and (iii) a ring with the density peak around the circumference of the inoculation drop (see Fig. S5). All three cases lead to the formation of a metabolic ring, and the final morphologies are qualitatively similar. The only remnants of the initial conditions are noticeable only at the very edge of the inoculation site. Thus, our simulations suggest that a broad high-density ring is primarily a product of the growth dynamics rather than the initial density pileup due to the capillary coffee-ring effect.

### Maintenance cost plays an important role in colony growth

It is well-known that cells spend energy not only for growth, but also for maintenance, e.g. repairing the cell wall, transcription, protein synthesis and degradation, sensing and navigating the environment, etc. The energetic cost of these routine tasks are completely neglected in the minimal models, which focus on growth only (some models allow for sporulation,^77^ but that is a distinct process). In contrast, FBA models typically include maintenance requirements, which leads to some unexpected results as we show below.

The *E.coli* core model contains a non-growth associated maintenance reactions that siphons away ATP with an experimentally estimated flux of 8.39 mmol/g·h^43,50^ (see Methods). As a result, growth stops in our simulations when the available resources are not sufficient to sustain a flux of ATP production in excess of 8.39 mmol/g·h.

To ascertain the ramifications of the maintenance cost, we varied the maintenance requirement from zero to about twice its actual value. Supplemental Fig. 7 illustrates the drop in the growth rate and biomass yield when the maintenance requirement is set to a non-zero value during growth in a well-mixed batch culture. In the case of colonies grown on plates, as simulated in COMETS, we similarly found that growth rate and yield also decreased significantly with the rise of the maintenance requirement (Fig. 7A).

**Figure 7.**
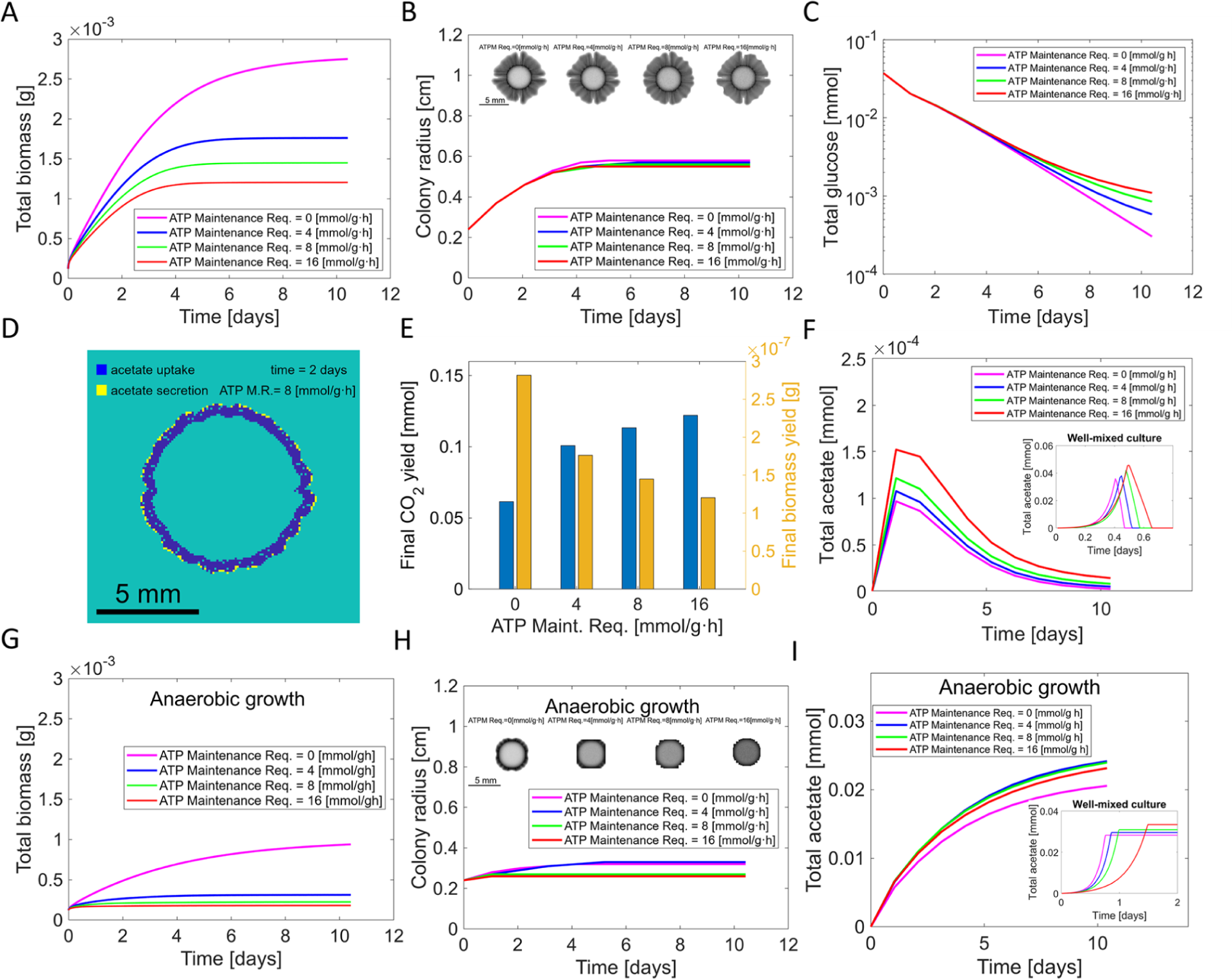
The colony biomass growth rate and yield decrease significantly with the rise of the ATP maintenance requirement, while the radius and morphology are only marginally affected. (A) The total colony biomass as a function of time for ATP maintenance requirement set to 0, 100%, and 200% of the actual maintenance cost in the *E. coli* model. (B) The colony morphology and average radius are not significantly affected by the maintenance requirement. (C) The time course of glucose depletion for different maintenance requirements is only marginally different at the final stages of the colony growth. (D) Acetate uptake and secretion display spatial heterogeneity within the colony. Acetate is secreted only by a thin sliver of rapidly growing cells at the colony edge. However, it is consumed deeper inside the colony, where glucose is depleted. (E) The final biomass and CO_2_ yield are the main terminal products of the metabolic reactions. The ratio of these two products depends strongly on the maintenance requirements. Higher maintenance results in more CO_2_ and less biomass. (F) Time progression of the total amount of acetate present in the entire (two dimensional) plate (main plot), compared to the well-mixed culture simulated with identical media, both in amount and concentration (inset). There is much less acetate accumulation in a colony compared to a well-mixed culture. In a colony, acetate is both produced and consumed at the same time, but in a liquid culture there is no acetate consumption until the cells run out of glucose and undergo a diauxic shift. The production and consumption of acetate are separated in space during colony growth, but they are separated in time for a well-mixed liquid culture. The low maximum acetate abundance in the spatially non-homogeneous case can be explained with the fact that different spatial regions undergo a diauxic shift from feeding on glucose to feeding on acetate at different times, creating a snapshot of the acetate exchange pattern in Fig. 7D. (G) Maintenance cost strongly affects biomass yield under anaerobic conditions. (H) The expansion rate is not as affected by the maintenance requirements as the biomass yield in (G). This is similar to the results in (A) and (B). (I) Under anaerobic conditions, *E. coli* cannot utilize acetate for energy. As a result, the acetate reaches similar levels under spatial and well-mixed conditions, but the temporal dynamics of the acetate accumulations are different reflecting the different rates of biomass growth: quasi-linear on agar plate and exponential in liquid culture.

Given this drastic dependence, we expected the colony morphology and the rate of radial colony expansion to also display a visible dependence on the maintenance flux. However, to our surprise, we saw that these quantities were hardly affected by maintenance (Fig. 7B). We trace these divergent responses to the different dynamics at the colony edge vs. the core of the colony.

At the edge, glucose concentration is the highest, and, therefore, the ATP production is much greater than the maintenance requirement. As a result, the growth and dispersal rates are only slightly reduced, and the rate of colony expansion remains virtually unchanged. The total biomass, however, also depends on the growth behind the colony edge, where the nutrient concentration is lower, and the maintenance requirement absorbs an appreciable fraction of the ATP produced thereby reducing both the growth rate and yield (see CO2 and biomass yield plots in Fig. 7E).

Consistent with this argument, we find that the glucose profiles are nearly the same across the maintenance costs (Fig. 7C), i.e. the nutrients get depleted at the same rate, and it is the conversion of nutrients into the biomass that explains the differences between the colonies in Fig. 7.

A deeper insight in the edge vs bulk dynamics can be gleaned from the spatial variations in the cellular metabolism (Fig. 7D). Near the edge of the colony, the cells are consuming glucose and secrete acetate. This metabolic pathway results in high ATP production rate and rapid growth. In contrast, the bulk of the colony has no access to glucose and is slowly growing on acetate. Thus there are not only quantitative differences in growth rates among the different regions of the colony, but also qualitative differences in their metabolism.^25^

COMETS offers the opportunity to explore the role of maintenance in a spatial setting under different metabolic constraints, e.g. in the absence of oxygen. From a direct comparison of Fig. 7G with Fig. 7A, we see that the colony growth in the absence of oxygen is also slowing down at higher maintenance requirements, as in the aerobic case. The time progression of the colony radius in Fig. 7H is not sensitive to the maintenance requirement, similar to the aerobic case in Fig 7B. The transition between fast and slow growth, however, occurs at a much lower maintenance cost when oxygen is not available. Specifically, we see a jump to a much lower growth rate already at 4 mmol/gh (Fig. 7G). This higher sensitivity to the maintenance cost is expected because ATP production is lower without oxygen. Finally, in the absence of oxygen acetate cannot be metabolized; as a result, its concentration raises and eventually saturates (Fig. 7I). The maintenance requirement at and above 4 mmol/gh does not affect either the growth rate or the time course of acetate production.

## Discussion

The plethora of shapes formed by bacterial colonies reflects the complexity of biological and physical phenomena at play in their formation. Many of these processes, including growth instability and sector formation, have been successfully studied using minimal single-nutrient models.^29,13,60,55^ The results of these minimal models, however, do not capture the full diversity of colony morphologies and cannot be directly applied to specific microbial strains growing in specific environments.

In this study we employed our software platform COMETS^54,27^ to show how the integration of physical processes and detailed metabolic information can recapitulate prior knowledge, provide new insights, and stimulate unexpected discoveries. What made this possible is the delicate balance between preserving computational efficiency and including detailed mechanistic descriptions of metabolism and cellular biophysics. We navigated this tradeoff by combining an efficient implementation of demographic fluctuations,^56^ a robust model of nonlinear biomass diffusion,^55^ and a genome-scale description of the bacterial metabolism^50^. Our results demonstrate that this level of detail is sufficient to capture many important biological processes, provide an accurate description of colony growth and even produce exceedingly realistic images of colonies themselves.

Systematic simulations of colony morphology as a function of nutrient (glucose) concentration and (nonlinear) biomass diffusivity reproduced the morphologies expected from previous experimental studies.^46,47^ Overall, our results were consistent with the theoretical expectation from minimal models that colony growth is largely controlled by a single parameter the product of biomass diffusivity and nutrient concentration. The detailed nature of our simulations, however, uncovered small, but noticeable deviations from this expectation, which are also present in the experimental data.^46,60^

Our simulations also shed light on how genetic drift and sector formation ^13,29,64^ depend on colony morphologies and growth conditions. Broadly speaking, we found that both branching and sector formation were delayed under favorable growth conditions, and, once formed, branches quickly lost diversity and became homozygous for one of the strains. This coupling between genetic demixing and colony morphology can be understood as follows. Local extinctions are common in spatially growing populations because local population sizes are small. Biomass diffusion, however, can restore the diversity by bringing a different strain from nearby spatial regions and thus delay genetic demixing. Branched morphologies interfere with the lateral transport of the biomass because each branch becomes an isolated subpopulation that cannot restore diversity by immigration. As a result, local extinctions quickly accumulate and each branch becomes homozygous for a particular strain.

One unexpected outcome of our simulations was the discovery of the metabolic ring. Typical minimal models aim to capture the growth at the front of the colony, so any structure in the colony bulk might be attributed to the artifacts of the model. The predictions of COMETS simulations should however be taken more seriously. Our simulations predicted a ring of high-density near the inoculation site, which matched closely our experimental observations. In previous experiments, this ring has been attributed to the coffee-stain effect,^66,68,69^ i.e. the accumulation of the cells near the edge of an evaporating drop right after inoculation. While this coffee-stain effect undoubtedly exists in bacterial colonies,^67^ it was perhaps premature to claim that it is the sole explanation of the high-density ring around the homeland because this ring appeared in our simulations which did not model the evaporation process.

We carried out a series of simulations to determine how initial conditions affect the high-density ring and found that the coffee-stain effect is responsible for only a tiny sliver of this extra density right near the inoculation site. The rest of the high-density ring is due to the accumulation of biomass during colony growth, which occurs with or without coffee-stain-like initial conditions. We also found that the metabolic ring becomes more prominent when biomass motility is suppressed, a prediction that was confirmed in our experiments with varying agar concentration (Fig. 6). We attribute this phenomenon to the following mechanism. Initially, the colony grows evenly, depleting the nutrients under the initial homeland. Once the nutrients are depleted in the center, they must diffuse from the outside, and, therefore, the growth is restricted to the edge of the colony leading to the formation of the metabolic ring. This transient dynamic stops once the colony starts to expand at an approximately constant velocity and no nutrients are available at the inoculation site. The duration of this transient increases with reduced biomass diffusivity resulting in the just-described dependence of the metabolic ring on the agar concentration.

The mechanistic model of *E. coli* metabolism that we used naturally included an energetic maintenance requirement and allowed us to investigate its effect on the dynamics and morphology of growing colonies. Although it is well-known that higher maintenance requirements lead to lower growth and lower yield in a well-mixed culture, it was not clear how this slower growth would manifest in a spatial context. To answer this question, we simulated colony growth for different values of the ATP maintenance flux in the metabolic network. The total biomass followed closely the predictions for the well-mixed populations: It grew slower and with lower yield. In contrast, the nutrient consumption rate and the spatial colonization rate, i.e. the increase in the colony radius over time, were virtually unaffected by the maintenance cost. This difference in behavior reflected the spatial separation of growth within the colony. The cells at edge received plentiful nutrients and grew at a maximal rate, which is only slightly affected by the maintenance cost. The cells deeper within the colony grew much slower and spent a significant proportion of their energy intake on maintenance. This spatial separation of the growth rates was also accompanied by changes in the metabolic fluxes within the cells, which further highlights the importance of both explicitly spatial modeling and the relevance of the metabolic costs.

The overall success of our simulation approach suggests that realistic *in silico* experiments are within reach not only for *E. coli* growing on a Petri dish, but for other bacteria growing in more complex environments. Here, we outlined an approach that combines genome-scale metabolic modeling with an appropriate physical model of biomass motility and demographic fluctuations. The ongoing efforts to construct metabolic networks for non-model organisms and to understand microbial motility in complex environments would enable application of our approach to practically relevant communities such as root-associated microbiota^22,20^ or biomedically relevant (e.g. dental) biofilms.^78–81^ For example, one could use COMETS to elucidate the synergistic role of diffusion in spatial microstructure and metabolic activity to shed light on the colonization of plant roots by rhizosphere bacteria.^20,21^ Such computational modeling can then further be employed to understand succession of aerobic and anaerobic communities, and even the consequences of blocking specific metabolic pathways in the community with drugs or genetic engineering.

Future development of our modeling approach certainly must address its current limitations and shortcomings. First, most communities of interest consist of many bacterial species, rather than the relatively simple community with two neutral strains that we studied here. We thus clearly see a need to improve automated procedures for model building and gap-filling from microbial genomes.^82,83^ Second, in our current model of biomass propagation, the two strains also have identical biophysical properties. In a system with non-identical species, the model of biomass propagation must take into account the differences in cell shape, cell-substrate and cell-cell interaction among different strains. Third, our modeling approach does not incorporate regulation, quorum sensing signals, or environment dependent change of metabolic objective. A future inclusion of these biological processes will greatly expand the possibility of *in silico* experiments in COMETS and will bring us closer to predictive ecosystem modeling.

## Acknowledgments

This work was partially supported by the U.S. Department of Energy, Office of Science, Office of Biological & Environmental Research through the Microbial Community Analysis and Functional Evaluation in Soils SFA Program (m-CAFEs) under contract number DE-AC02-05CH11231 to Lawrence Berkeley National Laboratory; the National Science Foundation (grants NSFOCE-BSF 1635070 and the NSF Center for Chemical Currencies of a Microbial Planet); the Human Frontiers Science Program (RGP0060/2021); the National Cancer Institute at the NIH (1R21CA279630-01); and the Boston University Biological Design Center Kilachand Multicellular Design Program. K.K. acknowledges funding from 1R01GM138530-01.

## Author contributions

Conceptualization, I.D., A.G., K.K., D.S.; Methodology, I.D., K.K.; Software, I.D.; Formal Analysis, I.D.; Investigation I.D., J.Z., M.O.; Resources, J.Z., M.O.; Data curation, I.D.; Writing - Original Draft, I.D.; Writing - Review and Editing, K.K., D.S.; Visualization, I.D.; Supervision, K.K., D.S.; Project Administration, K.K., D.S.; Funding Acquisition, K.K., D.S.

## Declaration of interests

The authors declare no competing interests.

## Inclusion and diversity statement

We support inclusive, diverse, and equitable conduct of research.

## Methods

### Key Resources Table

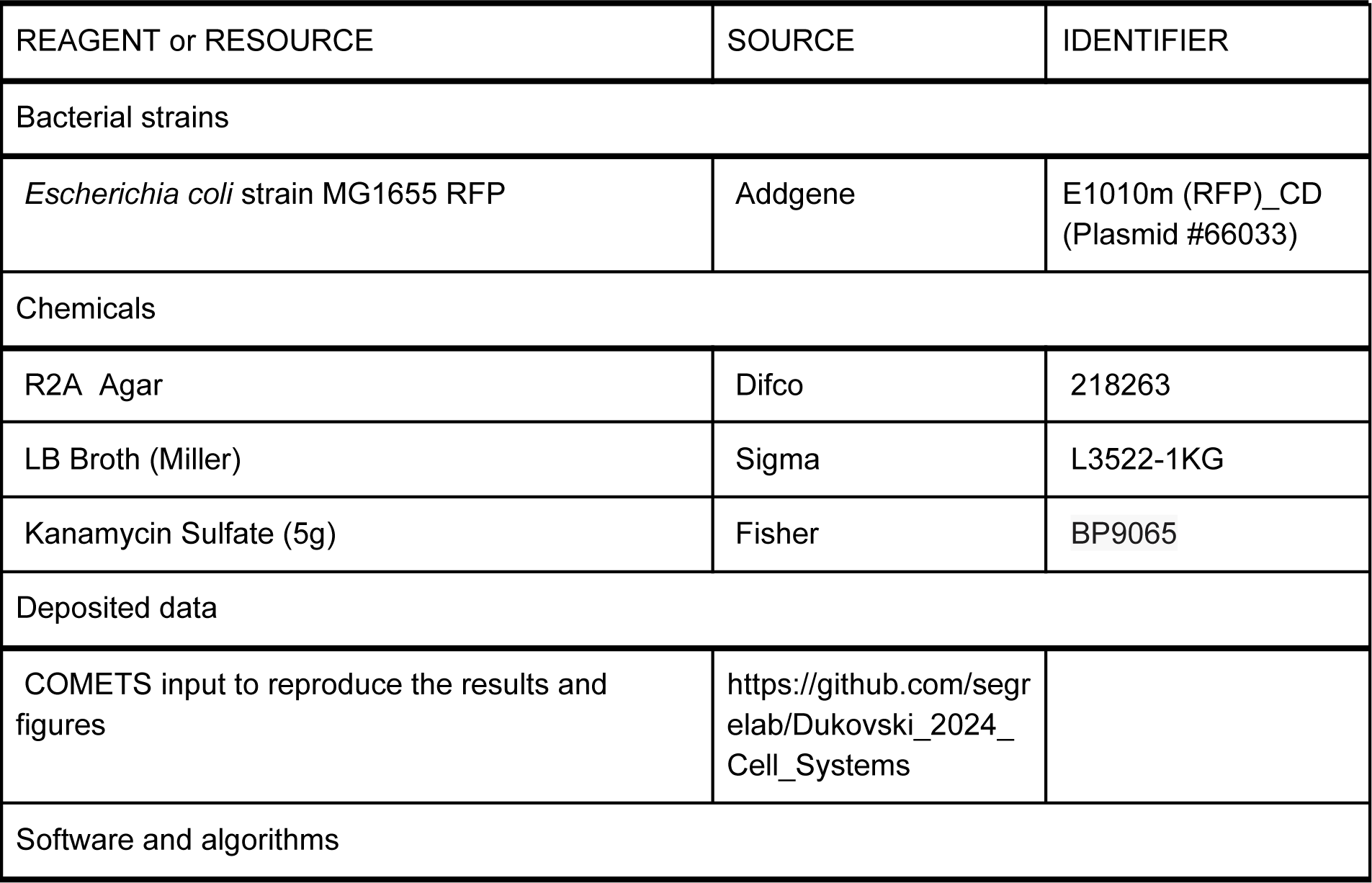
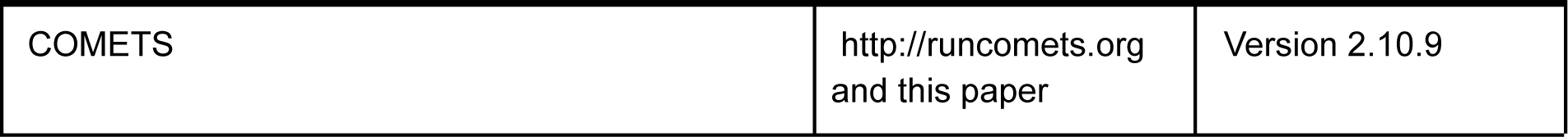

### Data and code availability

COMETS is a publicly available software. More information, user’s manual, including how it can be downloaded and installed are found at the COMETS website: http://runcomets.org COMETS is distributed under the GPL-3.0 license, and the source code can be downloaded from https://github.com/segrelab/comets. The computational input files, models, layouts and scripts to reproduce the results can be downloaded from https://github.com/segrelab/Dukovski_et_al_2024.

### Computational methods

In this study, we combined several modeling methodologies to simulate the growth and propagation of bacterial colonies. The three core modeling methodologies we utilized are: the spatio-temporal dynamic Flux Balance Analysis (dFBA), a model of cooperative spatial biomass propagation, and a model of population fluctuations or demographic noise. These three methodologies are synergistically implemented in our modeling platform for Computation of Ecosystems in Time and Space (COMETS).^54^

### Dynamic Flux Balance Analysis

The dynamic Flux Balance Analysis (dFBA)^57^ is a temporal extension of the Flux Balance Analysis (FBA)^84^ method for solving the mass balance requirements of a stoichiometric model of the metabolism (metabolic network) of an organism. Starting with the genome of an organism, a genome-scale reconstruction of the network of chemical reactions involved in the metabolism is produced. This metabolic reconstruction model is mathematically represented by the stoichiometric matrix *S* of the chemical reactions and the metabolites that form the metabolic network. Under the assumed constraints of mass conservation, steady state, and the maximum value of the nutrient uptake by the organism, the FBA method returns the values of the reaction fluxes through the metabolic network, including the rate of biomass production. In this study we calculated the upper limits for the uptake of each extracellular nutrient according to the Michaelis-Menten formula:

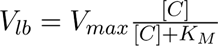

We obtained the values of the parameters *V_max_* and *K_M_* for the *E. coli* core model^50^ that we use in this study by comparing the simulated biomass growth curve of the model with the growth curve.^57^ We also reproduced the time dynamics of the secreted metabolites for this model with the same set of fitting parameters. The values that we used in this study are *V_max_* and 10.0*mmol*/*gh* and *K_M_* = 1.5 · 10^−5^*M*. We varied these values accordingly for the data presented in Figs. S2 and S3.

In Fig. 4a the growth rate obtained by the FBA calculations were slightly broadened (by about 3%) with a normal distribution. This broadening added a slight white Gaussian noise to the growth, with the aim to stimulate the branching transition of the growth front.

### Minimal metabolic model

In Fig. 2A and Fig. 2C we used a minimal model of metabolism, where a single nutrient is uptaken and transformed fully into biomass, with no secretion of any metabolites. The nutrient to biomass conversion coefficient was set to 1 g/mmol for Fig. 1A. For Fig. 1C, in order to match the biomass yield of the minimal model with the biomass yield of the *E. coli* core model, the coefficient for the nutrient was set to 12.79 g/mmol.

### Bacterial biomass propagation model

The bacterial biomass in COMETS is treated as a continuous variable, and biomass motility is described by non-linear diffusion in two spatial dimensions. The stochastic partial differential equation governing the biomass dynamics is:

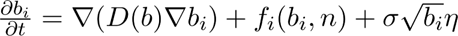

where on the left hand side is the time derivative of the local biomass *b_i_* of strain *i*. The first term on the right hand side is the spatial diffusion term. The diffusivity, which is assumed to be the same for all strains, depends only on the total biomass:

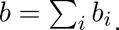

In its original form, the functional dependence of *D* on *b* is taken to be a power law

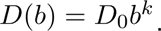

We used *K* = 1 throughout and further augmented this functional form with an additional cutoff at low growth rates. This was done to account for the arrest of growth and motility inside the colony, where the nutrients are depleted. We implemented the cutoff by multiplying the diffusivity with a Hill function of the relative portion of the biomass that has grown at each finite time step, with an adjustable cutoff *K_h_*:

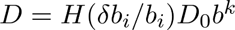

with *H* defined as

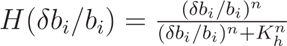

where *δb_i_* is the change of the biomass of species *i* grown during the time-step, and *n* is Hill exponent, which we took to be unity in all of our simulations. When the cutoff *K_h_* is set to zero, the entire biomass is propagating.

The local biomass growth term *f_i_*(*b_i_*, *n*) was computed via dFBA as described above. Note that *n* represents all metabolites in the local microenvironment. We assumed that the metabolites obey standard diffusion and took their diffusivity to be that of glucose in water with agar,^85^ *D_n_* = 6.0 · 10^−6^*cm*^2^/*s*.

The last term is the equation for biomass propagation is the demographic noise, which we describe in the next subsection.

### Demographic noise

We implemented the demographic noise following the method in reference [cite Munoz]. At each time step *t*, given the local nutrient contents, first we calculate the growth rate *μ* for the biomass using the FBA procedure described above. Then we calculate the local updated biomass *b̃*. To include a stochastic noise in the updated value of the biomass, we first sample a parameter *α* from the Poisson distribution:

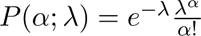

with *lambda* defined as

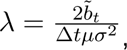

where *μ* is the growth rate obtained from the FBA optimization, and Δ*t* is the duration of the discrete time step. With *α* in hand, we sample *x* from the Gamma distribution:

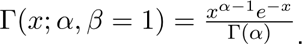

A preliminary (or proposed) stochastic value of the biomass is calculated as:

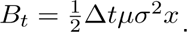

The next step is to reconcile the proposed value of the biomass with the FBA procedure. This extra step is necessary to ensure the stoichiometry of nutrient-to-biomass conversion, which in essence is the conservation of mass.

Given the proposed biomass *B_t_* in the current step, and the biomass in the previous time step *b*_*t*–Δ*t*_, we calculate new values for the maximum nutrient uptake rates (lower bound of the exchange reactions, since by arbitrary convention a negative exchange flux is uptake and positive flux is secretion), such that the newly FBA calculated biomass *b_i_* will be approximately close to the proposed stochastic one *B_i_*:

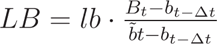

where *lb* is the lower bound of an exchange reaction calculated with the Michaelis-Menten dependence on the local nutrient concentration. Note that in the absence of stochasticity, the proposed biomass and the FBA calculated one would be the same.Thus, in the context of dFBA, demographic fluctuations are modeled as a density-dependent noise in the uptake rate.

After the noisy bounds on the uptake rates are determined, the FBA calculation is repeated yielding new values for the growth rate as well as metabolite consumptions and secretion rates. These are then used to determine the biomass and metabolite concentrations at this time step t. Note that, because the final updates are based on a dFBA calculation, the mass is strictly conserved.

Finally, we check that nutrient concentrations are positive and the biomass did not decrease. The latter would imply cell death, which is not included in our model. Such unphysical situations are rare, but unavoidable unless one deploys rather inefficient individual-based simulations. We remedy this by adjusting the lower bounds of the exchange reactions to avoid non-physical predictions and repeat the FBA calculation.^54^

### Simulations layouts

Three types of simulation layouts were used. In Fig. 2 and 5, and the corresponding supplemental figures, we created a 400×400 pixels grid simulating a 4×4 cm physical layout. The initial condition was a uniform fill of substrate nutrients consisting of a minimal medium of salts in unlimited supply (1000 mmol/pixel) and a finite amount of glucose. The boundary of the layout in Fig. 5 was kept at a constant glucose concentration throughout the simulation. The layout was initialized with a uniform biomass circle.

The layouts in Fig. 4 were 400×400 pixels grid simulating a 4×4 cm physical layout. The initial condition was a uniform fill of substrate nutrients consisting of a minimal medium of salts in unlimited supply (1000 mmol/pixel) and a finite amount of glucose. The layout was initialized with a linear inoculate of biomass along the entire bottom edge. The boundary at the top layout edge was kept at a constant glucose concentration throughout the simulation. This setup ensured the propagation of the growth front in only one direction.

The layout of Figures 3, 6 and 7, and the corresponding supplemental figures, were designed to approximate a 6-well plate. We used a 350×350 pixels grid to model a 3.5×3.5 cm spatial domain. The simulated plates were initialized with a minimal medium with glucose, and nutrients were allowed to deplete over time; we used reflecting boundary conditions. The initial distribution of the biomass was a uniformly populated circle.

The simulations were executed on the BU SCC cluster. A typical run used 8 CPU cores to solve the FBA problem at several grid points in parallel, and lasted for several hours. All simulation layouts are available for download at: https://github.com/segrelab/Dukovski_2024_Cell_Systems.

### Experimental methods

We grew colonies of *E. coli* strain MG1655, with added RFP plasmid Addgene E1010m (RFP)_CD.^86^ The initial streaking was done on LB agar substrate with added Kanamycin. Batch culture was grown in LB, diluted and inoculated 3μL on agar 6-well plates. The agar plates’ substrate was R2A with added 0.5 g/L glucose. Six concentrations of agar were used for each plate: 2, 4, 7, 10, 15, 25 g/L. Three replicas of the plates were held for 7 days at 30°C. Images were taken daily with 4x magnification Nikon Eclipse Ti2 microscope. One control replica was not taken out of the incubator and not imaged, to assure that the daily imaging did not disrupt the growth of the colonies. The images were post-processed with the NIC Nikon software. The photo images were taken with a Nikon D850 camera, 50mm+extension tube objective.

## Notes

https://github.com/segrelab/Dukovski_et_al_2024

### Competing Interest Statement

The authors have declared no competing interest.

