## Supplemental Information for "Biophysical metabolic modeling of complex bacterial colony morphology"

### Supplemental Video Title and Legend

**Supplemental Video 1. Related to Figure 7. A growth sequence demonstrating the formation of the metabolic ring.** The top sequence is a simulated colony viewed from above. The bottom sequence shows the dynamics of estimated colony heights based on the biomass content in each pixel of the simulation grid, assuming that the biomass density is the same as that of water. The formation of the ring is at the onset of the growth of the biomass at the edges of the homeland (initial inoculum).

### Supplemental Figures

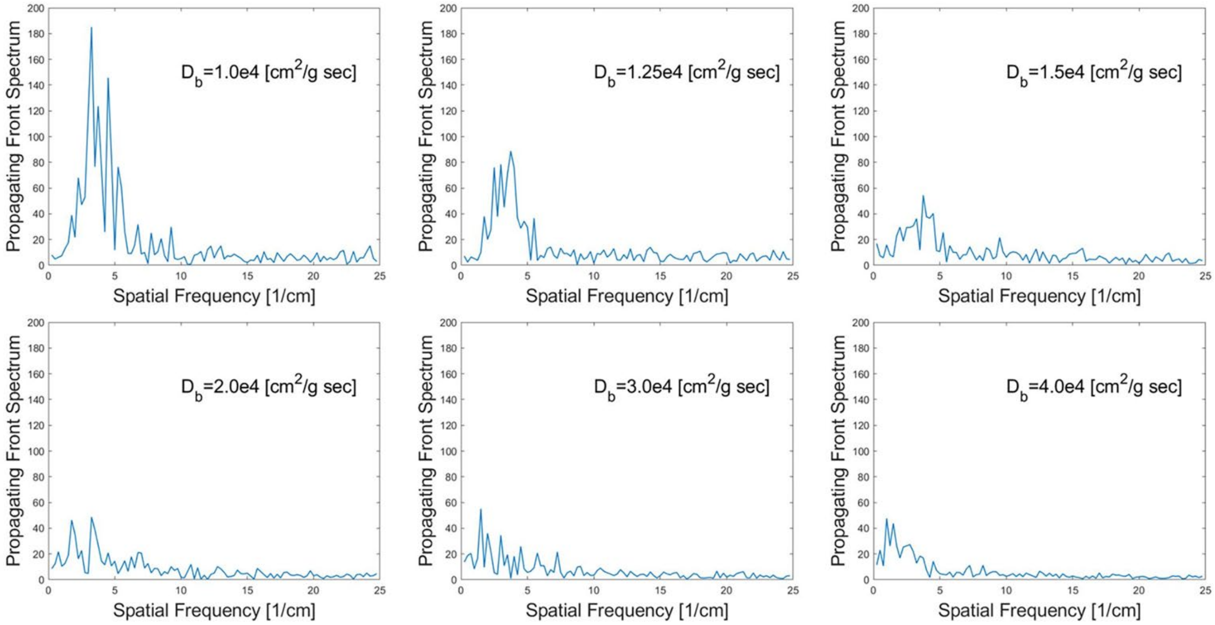


**Supplemental Figure 1. Related to Figure 4.** Fourier transform of the linear colony growth front for several values of the biomass diffusivity prefactor. The peak of the transform relates to the depth and characteristic length of the branching profile of the front. The Furier transform peak is higher for lower values of the biomass diffusivity, and levels down to an approximate constant value for the lowest three values. We define the transition point from branched to smooth morphology as the transition from decaying to constant peaks.

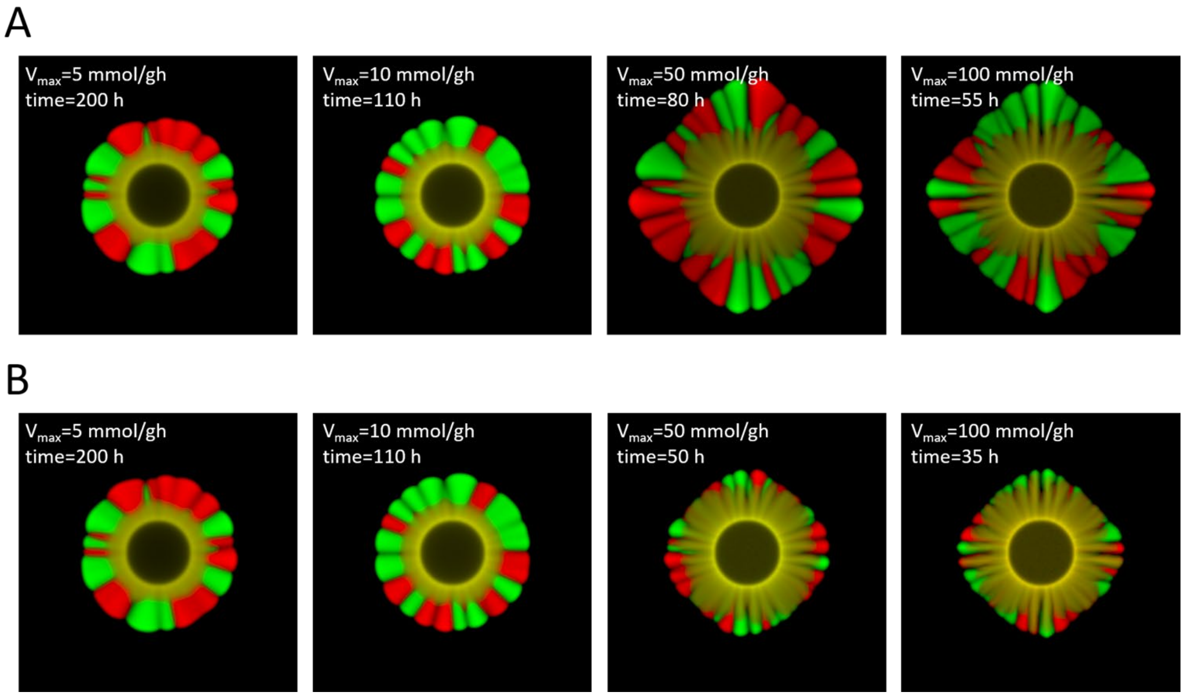


**Supplemental Figure 2. Related to Figure 5.** The sensitivity of the colony morphology to the change of the maximum nutrients uptake rate parameter of the genome-scale model.

(A) Images of the colonies grown with several values of the maximum uptake parameter, taken at the time points where the biomass in each case is the same.

(B) Images of the colonies grown at the same conditions as (A), but taken at the same approximate colony radius.

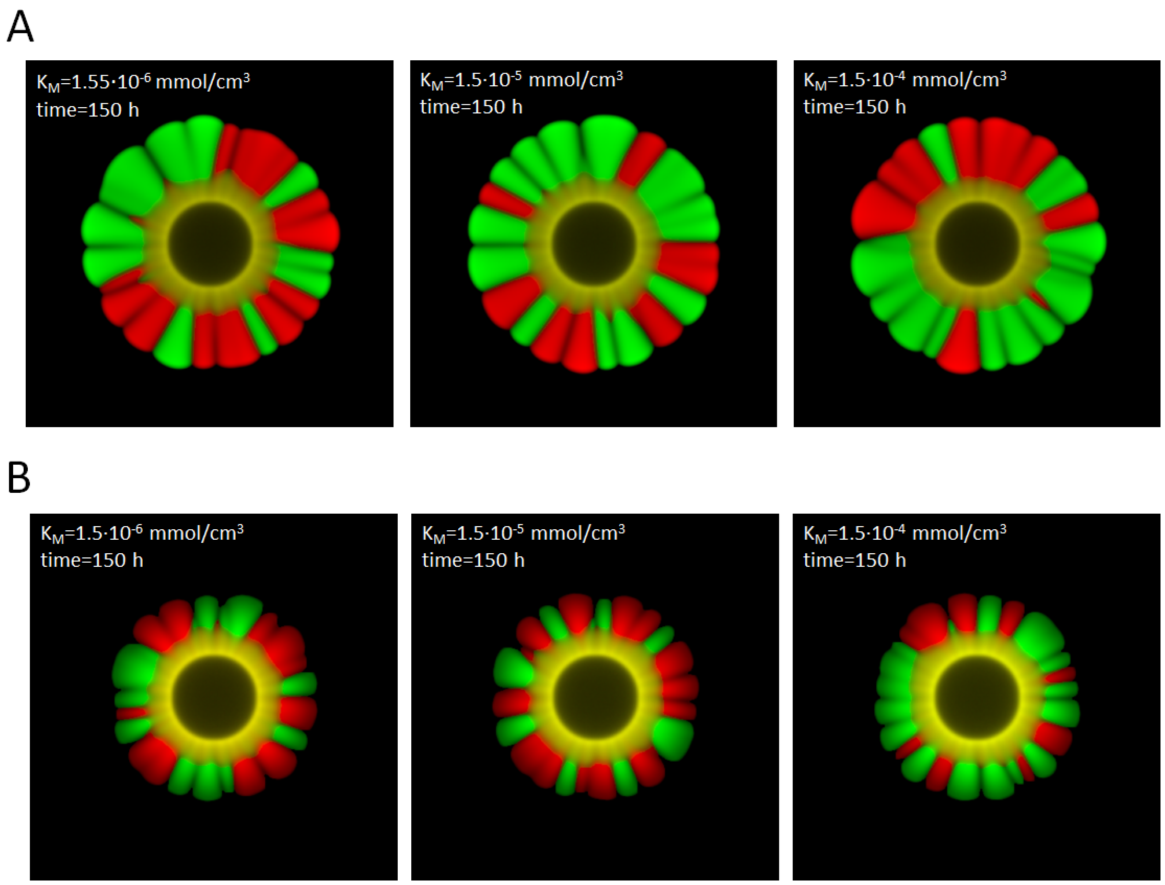


**Supplemental Figure 3. Related to Figure 5.** The morphology of the colony is not sensitive to the value of the Michaelis constant in the Michaelis-Menten uptake function of the genome-scale metabolic model. The value used in this study is the middle one, $K_M$=1.5·10$^{-5}$ mmol/cm$^3$.

(A) The colonies are grown with constant concentration of nutrients at the boundary. The nutrients are not depleted, the colony does not seize growing.

(B) The nutrients deplete from their initial uniform abundances, and are not replenished. he colony stops growing when the nutrients are fully depleted.

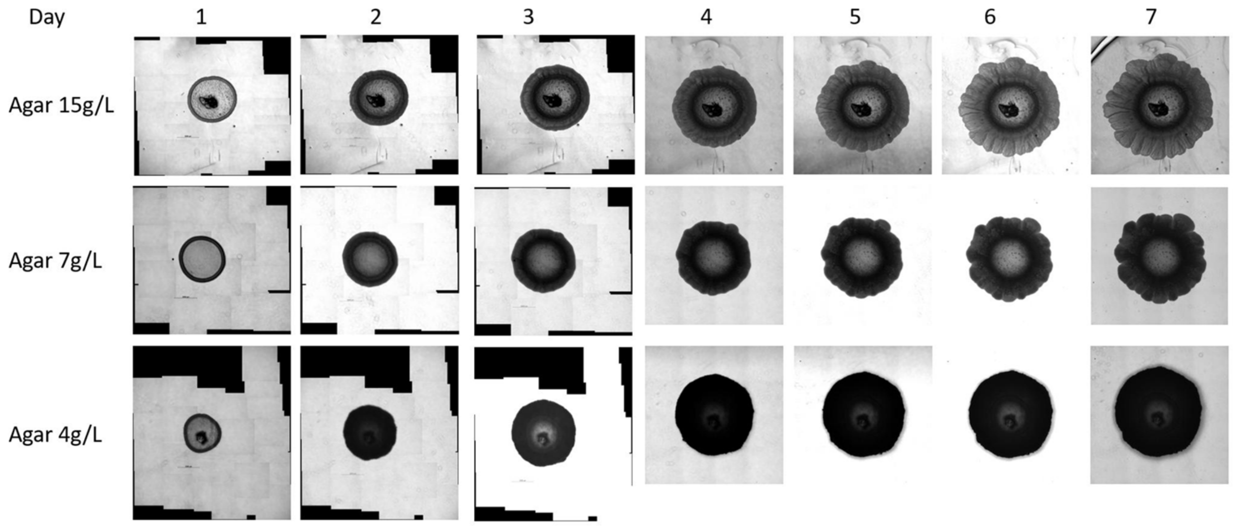


**Supplemental Figure 4. Related to Figure 6.** Experimentally grown E. coli colonies on substrates with three different agar concentrations. The initial stage of the colony shows the capillary coffee-stain ring. The capillary ring is the same for all three agar hardnesses. The colony on the hardest agar, however, develops most prominent metabolic ring over time, as a ring shaped plato distinguishable from the rest of the colony.

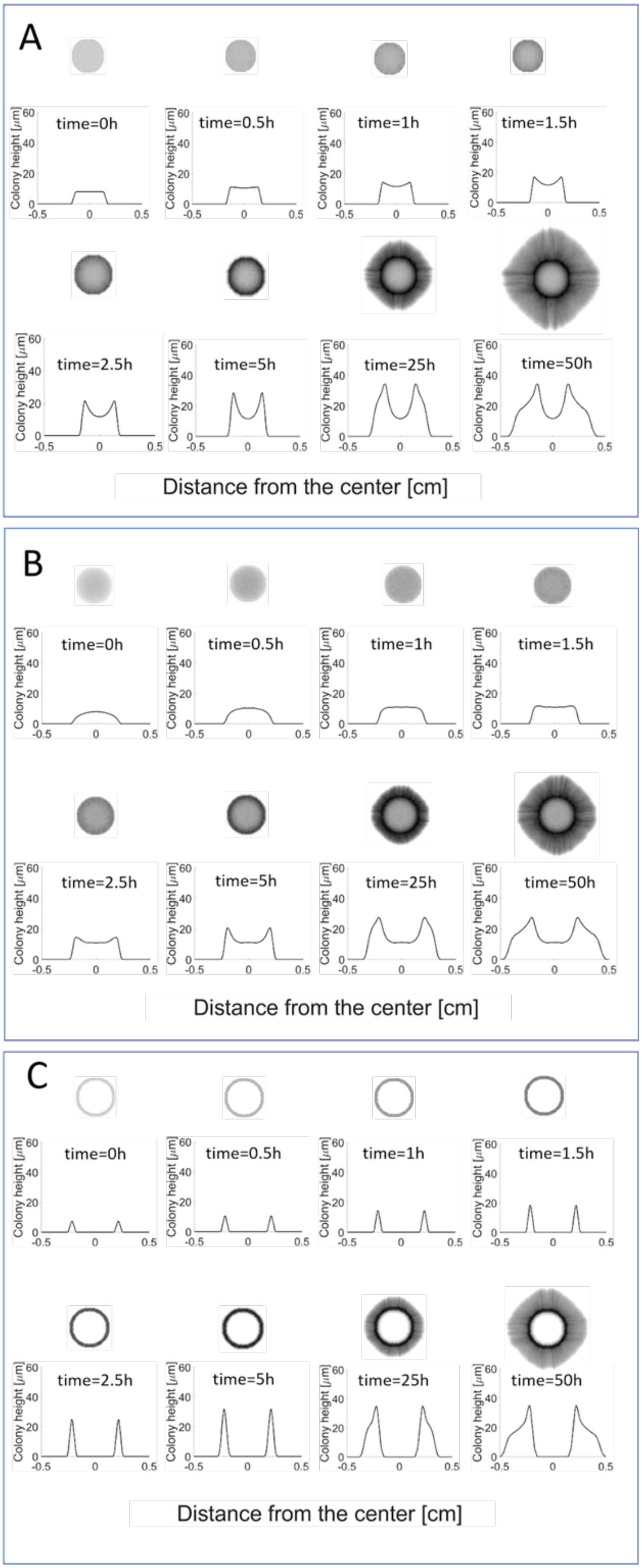


**Supplemental Figure 5. Related to Figure 6.** The metabolic ring is formed regardless of the shape of the initial inoculum:
(A) flat "pancake",
(B) hemispherical dome,
(C) simulated physical coffee-stain ring.

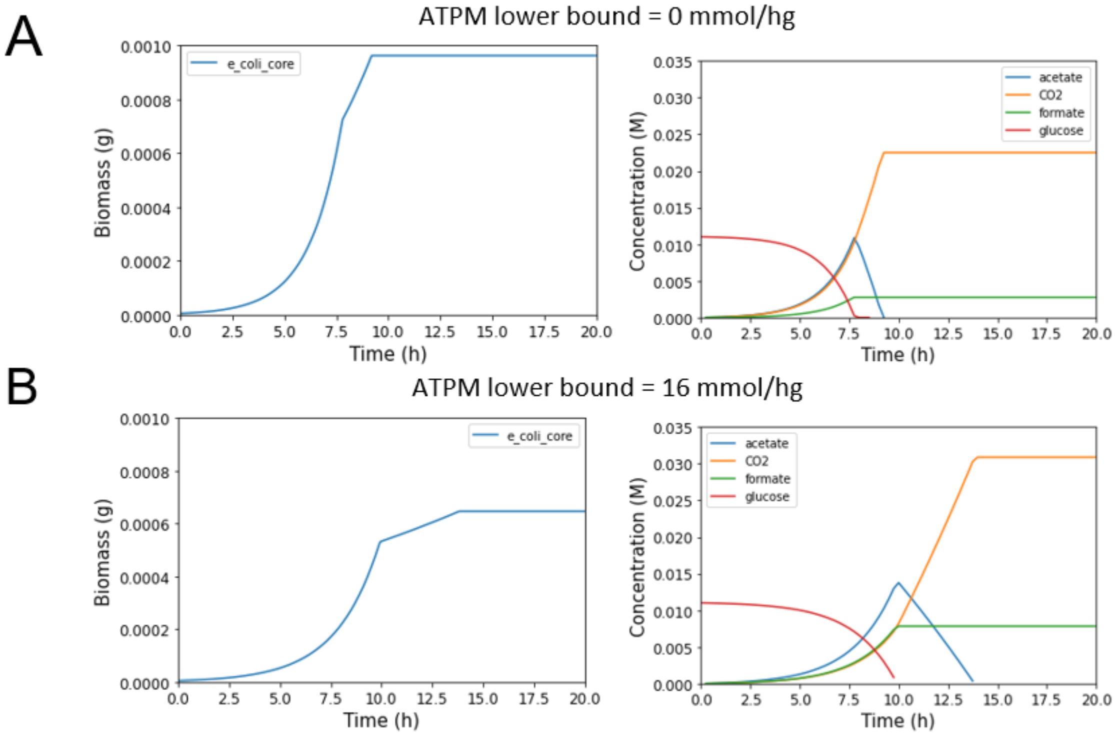


**Supplemental Figure 6. Related to Figure 7.** Simulations of E. coli well-mixed culture with (A) zero and (B) 16 mmol/g·h ATP maintenance requirement levels. Both the biomass growth rate and final biomass yield are affected by the requirement change, with lower growth and lower yield for higher ATP maintenance requirement. The maximum glucose and the product of the metabolism, most notably acetate, concentrations are of comparable order of magnitude, in contrast to the significant difference in the spatially structured colonies in Fig. 7.
